# Rescue from pyridostatin-stabilized telomeric G-quadruplexes by DNA cleavage and microhomology-mediated end joining

**DOI:** 10.1101/2025.11.25.690390

**Authors:** Samah Matmati, Satyajeet Rao, Joachim Lingner

## Abstract

G-quadruplexes are four-stranded nucleic acid structures that are formed by guanine rich sequences. They have been implicated to interfere with telomere maintenance, but their effects are not well understood at a molecular level. Here, we treat human cells with the G-quadruplex stabilizer and anti-cancer drug-candidate pyridostatin and characterize its effects on telomere replication and maintenance. We demonstrate that pyridostatin interferes with lagging strand DNA synthesis during semiconservative replication of telomeres. Pyridostatin impedes replication fork progression and triggers telomere cleavage events involving the Flap-endonuclease FEN1 and the nuclease-helicase DNA2. Telomeric protein composition analysis reveals pyridostatin-induced association of LIG3 with telomeres. This enzyme is involved in repair of DNA double strand breaks by microhomology-mediated end joining (MMEJ). Depletion of MMEJ-factors or their chemical inhibition enhances telomere loss events in pyridostatin-treated cells. Altogether, our data reveal a three-step mechanism in which pyridostatin-stabilized G-quadruplexes inhibit replication of lagging strand telomeres, which is overcome by FEN1/DNA2-mediated DNA cleavage and MMEJ-mediated DNA repair. The synthetic effects of pyridostatin and MMEJ inhibition on telomere maintenance may become exploitable in future anti-cancer therapies.

## Introduction

Telomeric G-rich sequences have a high propensity to adopt a stable secondary structure known as G-quadruplex (G4) (Parkinson *et al*, 2002; Wang & Patel, 1993). In G4 structures, four guanines form a planar array stabilized by Hoogsteen base-pairing, and multiple G-quartets stack on each other to form a stable, compact structure (Parkinson *et al*, 2002). Telomeric G4 formation may occur in the G-rich telomeric strand when DNA becomes transiently single stranded during DNA replication, at the single stranded G-rich 3’ overhang or at telomeric R-loop structures in which the G-rich DNA strand is displaced (Varshney *et al*, 2020). The conservation of G-rich sequences at telomeres during evolution has promoted speculations that G-quadruplexes at telomeres must provide some function (Rhodes & Lipps, 2015). However, it is better established that excessive or unresolved G4 may cause replication stress and damage at telomeres. Indeed, numerous helicases (Mendoza *et al*, 2016) including RTEL1 (Vannier *et al*, 2012; Deng *et al*, 2013), BLM (Sfeir *et al*, 2009; Vannier *et al*, 2012; Yang *et al*, 2020; Drosopoulos *et al*, 2015), WRN (Crabbe *et al*, 2004), FANCJ (Castillo Bosch *et al*, 2014) and PIF1 (Paeschke *et al*, 2013) have been identified that can resolve G4 structures apparently in a non-redundant fashion as deficiencies of individual helicases lead to severe disease due to genome instability. In addition, the single strand DNA binding proteins POT1 (Zaug *et al*, 2005), CST (Zhang *et al*, 2019) and RPA (Maestroni *et al*, 2017) counteract G4 possibly by binding and thus trapping the unfolded DNA.

Pyridostatin (PDS) is highly selective G4-binder that was designed to target polymorphic G4 structures (Rodriguez *et al*, 2008; Müller *et al*, 2012). It recognizes G-tetrads through π-π stacking interactions. G-tetrad affinity of PDS is further increased through hydrogen bonding and electrostatic interactions (Liu *et al*, 2022). PDS has been shown to induce DNA damage foci at telomeres and at G-rich regions elsewhere in the genome (Rodriguez *et al*, 2012). Furthermore, PDS reduces the viability of cancer cells that are deficient in homology directed repair elevating the levels of replication problems and DNA damage (Zimmer *et al*, 2016; Groelly *et al*, 2022; Keahi *et al*, 2025). G4-stabilizing drugs can also inhibit telomerase as well as inhibit telomeric C-strand DNA synthesis post elongation of the G-strand by telomerase (De Cian *et al*, 2007; Johnson *et al*, 2024; Bryan, 2020).

In this study, we demonstrate that PDS impairs DNA replication fork progression at telomeres during semiconservative DNA replication interfering with lagging strand DNA synthesis. Paused replication forks at telomeres are cleaved in a FEN1/DNA2-dependent manner and repaired by MMEJ thus overcoming PDS-induced replication stress. Upon inhibition of MMEJ, PDS-treated cells suffer from severe telomere damage due to sudden telomere cleavage events. Thus, MMEJ is crucial for telomere integrity and telomere maintenance under G4-stabilized conditions.

## Results

### PDS induces telomere fragility and telomere cleavage at lagging strand telomeres

Replication stress at telomeres has been demonstrated to induce telomere fragility (Sfeir *et al*, 2009). It is characterized by smeared or doubled telomeric FISH signals on metaphase chromosomes. To investigate the effects of PDS on telomere replication we measured telomere fragility in HeLa cells treated for 24 hours with 0.12 µM PDS (Fig. 1A, B). Telomere fragility increased from approximately 3% in untreated cells to an average of 9% in PDS-treated cells, indicating interference with telomere replication. At the same time, a low but significant number of chromosome ends with detached telomeres (so-called outsiders (Majerska *et al*, 2018)) accumulated indicating that G4-stabilization by PDS induces telomeric breakage events (Fig. 1A,C). Of note, the loss of even a single telomere in a cell may have lethal consequences as it can promote chromosome rearrangements, degradation and loss events. Thus, the observed frequency (0-8%; average of 1%) of outsiders upon PDS treatment could have detrimental consequences.

**Figure 1.**
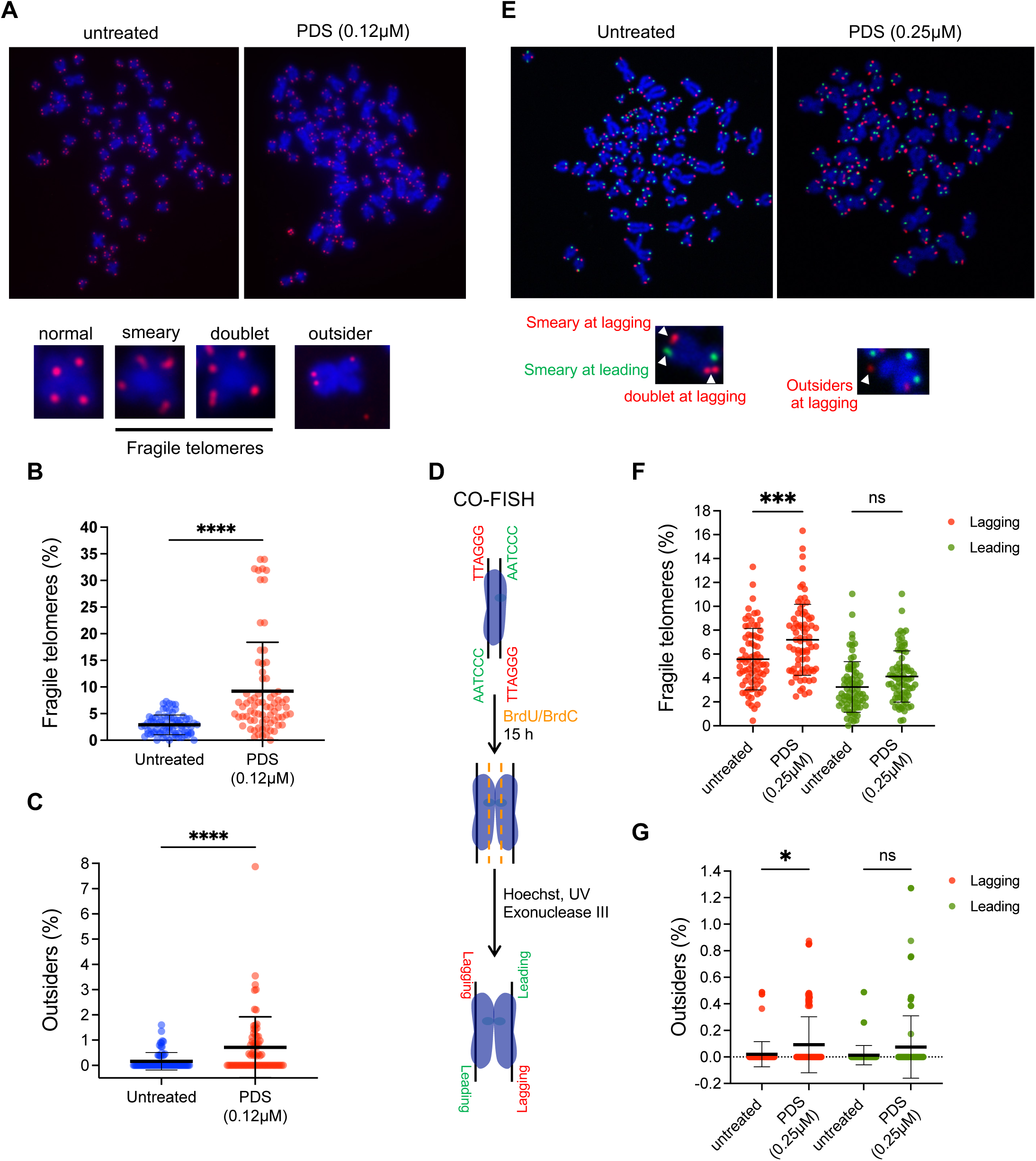
PDS induces telomeres fragility at lagging strand telomeres. (**A**) Representative telomeric FISH images of metaphase chromosome spreads of HeLa cells (with 10 kb average telomere length) stained with telomeric Cy3-[CCCTAA]_3_ FISH probe (red) and DAPI (Blue). Untreated (upper left) and PDS treated (upper right) samples are shown. Examples of normal, fragile (smeary or doublet telomeric signals) or outsider (telomeric FISH signal detached from a chromosome arm) telomeres are shown on the bottom. (**B**) Quantification of fragile and (**C**) outsider telomeres in untreated cells and upon treatment with 0.12 µM PDS for 24 h. 75 metaphase spreads were analyzed per condition, across three independent biological replicates. Horizontal lines and error bars represent mean ± s.d. P-values are calculated from Mann Whitney test (\*\*\*\**P* < 0.0001). (**D**) Schematic of CO-FISH experiment in HeLa cells (with more than 30 kb average telomere length). Mitotic cells are harvested after exposure to BrdU/BrdC for 15 h. Newly synthesized DNA strands containing BrdU/BrdC are selectively degraded upon Hoechst staining, UV irradiation and exonuclease III digestion, leaving only the parental strands intact. The remaining parental DNA strands are detected upon hybridization with strand-specific probes. The black lines indicate the parental DNA strands, the dashed orange lines the newly synthesized DNA strands. The chromosome ends replicated by lagging or leading strand synthesis are in red or green, respectively. (**E**) Representative telomeric CO-FISH images of metaphase chromosome spreads of HeLa cells (with >30kb telomeres) untreated or treated with 0.25 µM PDS for 24 h and stained with TYE563-TeloC LNA probe (red), FAM-TeloG LNA probe (green) and DAPI (blue). White arrowheads (bottom) indicate strand specific fragile telomeres or outsider telomeres. (**F**) Quantification of fragile telomeres and (**G**) outsiders at lagging or leading strand telomeres. 75 metaphase spreads were analyzed per condition, across three independent biological replicates. P-values are from ordinary one-way ANOVA (\*\*\**P* = 0.0006, \**P* = 0.0127).

To assess whether there is any strand-specificity of telomere fragility upon PDS treatment, we performed telomeric chromosome-orientation FISH (CO-FISH) (Bailey *et al*, 1996). In brief, HeLa cells were incubated with BrdU and BrdC for 15 hours for incorporation of the modified nucleotides into the newly synthesized daughter strands (Fig. 1D). After spreading metaphase chromosomes on slides and staining with Hoechst dye, slides were UV irradiated, which preferentially damages the newly synthesized DNA strands containing the modified nucleotides. Following exonuclease III treatment, which degraded the nascent gap-containing DNA, the remaining G-rich parental strands were hybridized with 6-FAM-labeled LNA probes, while the parental C-rich strands were detected by hybridization with TYE563-labeled LNA probes (Fig. 1E). For optimal staining and imaging conditions, HeLa cells with >30 kb long telomeres (Cristofari & Lingner, 2006) were used to perform CO-FISH and PDS concentration was adjusted to 0.25 µM (Fig. EV1). As expected, G4 structures accumulated at telomeres in PDS-treated cells as assessed by a proximity ligation assay (PLA) using antibodies recognizing G4-structures and the telomeric shelterin component TRF2 (Fig. EV1A, B). Telomere fragility in HeLa cells with >30 kb long telomeres increased from an average of 12% to 21% upon PDS treatment (Fig. EV1C) and outsider telomeres also accumulated as already seen in HeLa cells with 10 kb telomeres, albeit at lower levels (Fig. EV1D). CO-FISH analysis of telomere fragility in PDS-treated cells revealed a significant increase in fragility at telomeres replicated by lagging strand synthesis whereas at leading strand telomeres the slight increase in fragility did not reach statistical significance (Fig. 1E, F). Outsider telomere frequency also increased significantly only at lagging strand telomeres (Fig. 1G). The specific effects seen at lagging strand telomeres are consistent with the fact that the telomeric G-rich strand used as DNA template is partially single stranded during the discontinuous DNA synthesis, facilitating the formation of G4s stabilized by PDS at lagging strand telomeres. At leading strand telomeres, on the other hand, the G-rich strand is newly synthesized remaining fully base-paired with the parental C-rich template strand. Thus, PDS may not efficiently induce G4 structures at leading strand telomeres during replication.

### PDS impairs replication fork progression at telomeres

While telomere fragility is a well-established readout for telomere replication problems, the assay is indirect and does not offer a high enough resolution to reveal the molecular structure of fragile telomeres. We set out to confirm replication stress at telomeres upon PDS treatment, utilizing complementary assays. Activation of the ATR checkpoint kinase at stressed replication forks leads to phosphorylation of RPA32 at Ser33 (P-Ser33) (Vassin *et al*, 2009) providing an independent assay for replication stress. We detected P-Ser33 RPA32 foci by immunofluorescence and quantified their colocalization with telomeric DNA detected by FISH in non-synchronized cells (Fig. 2A, B). PDS treatment induced a significant increase of P-Ser33 RPA32 foci at telomeres indicating increased replication stress.

**Figure 2.**
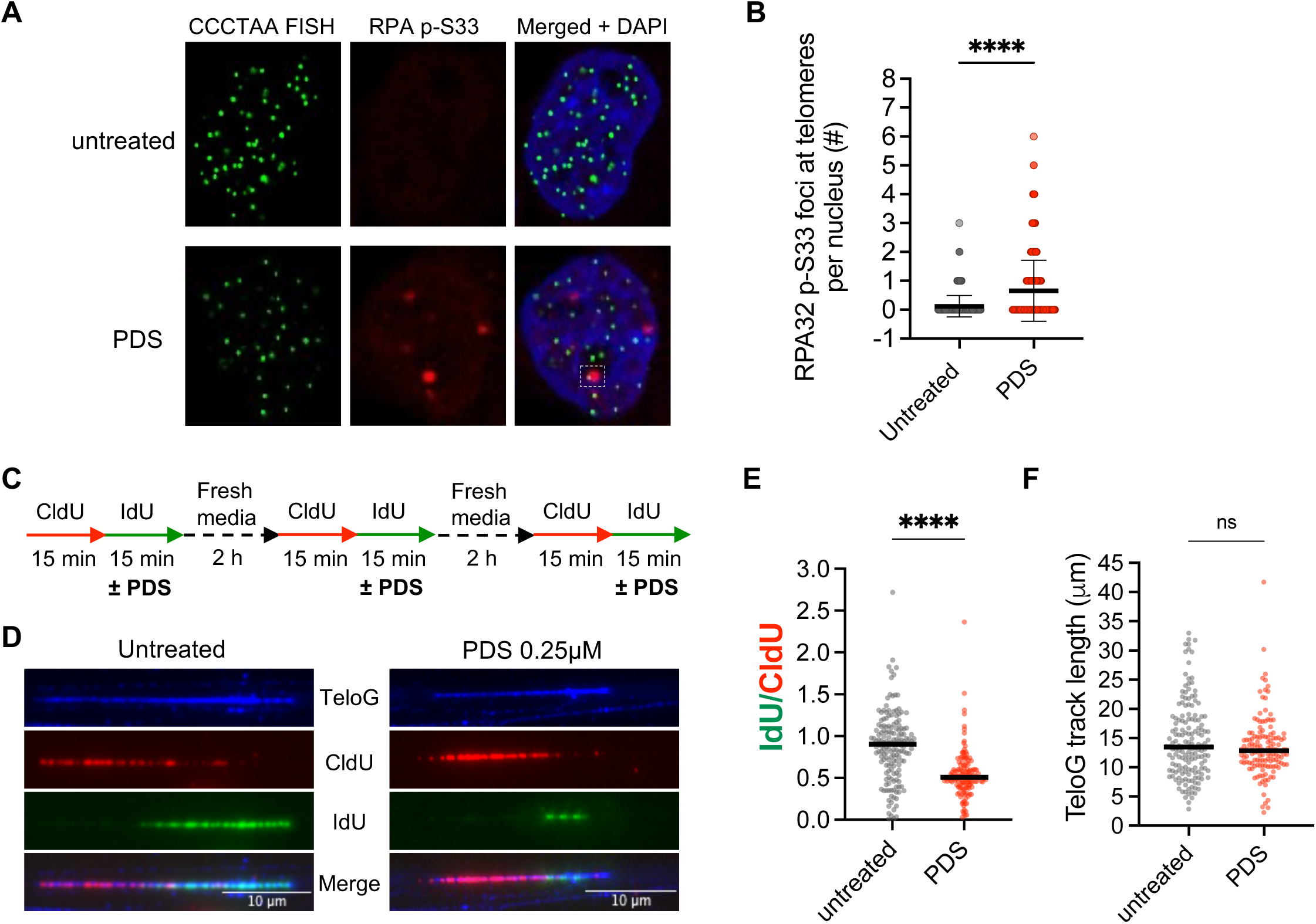
PDS impairs replication fork progression at telomeres. (**A**) Representative images for detection of phosphorylated RPA32 (P-Ser33) at telomeres. Immunofluorescence (IF) for RPA32 (P-Ser33) (in red) was combined with telomeric Cy3-[CCCTAA]_3_ FISH (in green) and DAPI staining (in blue). The dashed white square indicates a co-localized RPA32 (P-Ser33) focus with a telomere. (**B**) Quantification of RPA32 (P-Ser33) foci co-localizing with telomeres per nucleus. Scatter plot with bars indicating the mean ± s.d. from three independent biological replicates. P-value was calculated from Mann Whitney test (****P < 0.0001). (**C**) Workflow for DNA fiber labeling performed in Hela cells (with >30kb telomeres). Cells were pulsed with thymidine analogs (CldU and IdU) for 15min each. 0.25 µM PDS was added together with IdU. To increase the number of telomeric DNA molecules stained with the thymidine analogs, the labelling was repeated three times. (**D**) Representative images of replicated telomeric tracks with telomeric DNA (blue), CldU (red) and IdU (green). (**E**) Scatter plot of the IdU/CldU tract length ratios with bars indicating the median. At least 150 replication tracts were scored in three independent experiments for each condition. The P-value was calculated from the Mann Whitney test (****P < 0.0001). (**F**) Scatter plot of telomeric track length detected with TeloG probe in untreated or with 0.25 µM PDS treated cells with bars indicating the median. The P-value was calculated from the Mann Whitney test (ns = 0.3344).

To directly compare DNA replication fork progression at telomeres in presence and absence of PDS, we employed single molecule analysis of replicated telomeric DNA (SMARD) (Norio *et al*, 2005). In brief, replicating DNA in HeLa cells with >30 kb telomeres was sequentially labeled three consecutive times during 15 minutes with 5-chloro-2’-deoxyuridine (CldU) and 5-iodo-2’-deoxyuridine (IdU) (Fig. 2C). PDS was added concomitantly with IdU to check its effects on fork progression. Telomeric DNA was enriched by cleaving genomic DNA into small fragments with four base-pair cutting restriction enzymes (HinfI and RsaI) leaving the telomeric DNA intact. The telomeric DNA being much larger than the bulk of restricted genomic DNA was separated by gel electrophoresis and analyzed under the microscope detecting telomeric DNA by FISH and newly synthesized DNA by staining with anti-CldU and anti-IdU antibodies (Fig. 2D, E). The CldU and IdU-tract lengths gave a measure for replication speed of telomeric DNA. In the absence of PDS, CldU and IdU were incorporated at a similar rate (Fig. 2D, E). Very strikingly, however, PDS treatment during the IdU pulse caused a nearly two-fold reduction in IdU-fiber length demonstrating the strong interference of PDS with telomere replication. Overall telomere length as assessed by measuring the telomeric G-strand by FISH was not affected by the short-term treatment with PDS (Fig. 2F).

### PDS treatment triggers accumulation of LIG3 at telomeres

In order to elucidate the cellular response on PDS treatment at telomeres, we compared the telomeric protein composition of untreated and PDS-treated cells by 2-step QTIP (Glousker *et al*, 2020); Fig. 3A). For this analysis, we used HEK293E cells expressing flag-tagged TRF1 and TRF2 (Lin *et al*, 2021) that could be grown in suspension at large scale. The cells were treated for 24 hours with 15 µM PDS resulting in telomere fragility levels comparable to those in HeLa cells treated with 0.12 µM PDS (Fig. EV2A). Crosslinked and sonicated telomeric chromatin was purified from control and PDS-treated HEK293E cells first with bead-associated anti-flag antibodies that pulled down telomeric chromatin via flag-tagged TRF1 and TRF2 proteins. Upon elution of bound chromatin with excess of flag-peptide, telomeric chromatin was further enriched using affinity-purified polyclonal antibodies against TRF1 and TRF2 (Fig. 3A). The recovery of telomeric DNA was 12% and the enrichment of telomeric DNA over Alu-repeat DNA was roughly 2,000-fold (Fig. 3A, B, EV2B) indicating high purity of telomeric chromatin. TMT-labeling and analysis by mass spectrometry of untreated and PDS-treated telomeric chromatin allowed identification of 501 proteins (Fig. EV2C). The telomeric protein composition of PDS-treated cells was markedly altered (Fig. 3C, D; Table EV1). The telomeric shelterin proteins TPP1, RAP1, TIN2 and POT1 were slightly enriched with respect to TRF1 and TRF2. RECQ5 and TOP1 which counteract replication stress were also more abundant at PDS-damaged telomeres and several heterochromatin proteins CBX5 and LRWD1 were enriched. Most significantly, LIG3 was strongly enriched in PDS-treated telomeric chromatin.

**Figure 3.**
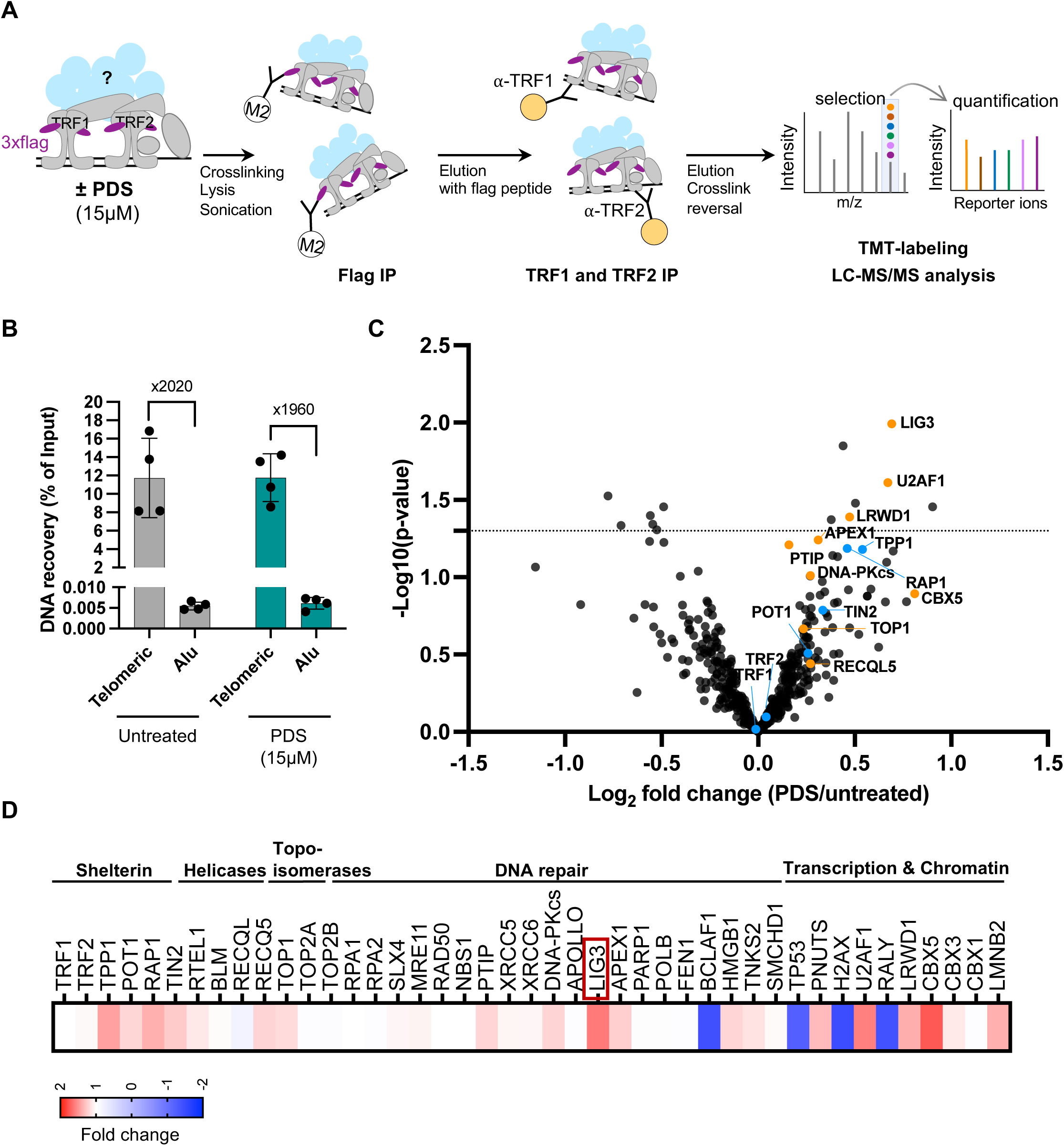
2-step QTIP reveals PDS triggered accumulation of LIG3 at telomeres. (**A**) Schematic of 2-step QTIP. Isolation and analysis of telomeric proteome was done using HEK293E cells expressing 3xflag-tagged TRF1 and TRF2 alleles. Cells were grown in suspension and treated with 15 µM PDS for 24 h. After crosslinking, lysis and sonication, telomeric chromatin fragments were first immunopurified using anti-flag M2 beads. Upon elution with flag peptide, telomeric chromatin was further enriched with anti-TRF1 and TRF2 antibodies. Telomeric chromatin was then analyzed by mass spectrometry (MS) using tandem mass tag reagents (TMT) and Liquid Chromatography-MS/MS (LC-MS/MS). (**B**) Quantification of recovered telomeric DNA (% of input) upon 2-step QTIP and fold enrichment of telomeric DNA over Alu repeat DNA (based on dot blot analyses). Bars represent data from four independent experiments ± s.d. (**C**) Volcano plot representing differential protein expression in untreated versus PDS-treated samples. P-value were determined using LIMMA followed by a Benjamini-Hochberg multiple-testing correction method. Proteins above the black dashed line are P < 0.05. Shelterin components are indicated with blue dots. Other relevant candidates enriched upon PDS treatment are indicated with orange dots. (**D**) Heat map representing changes in levels of selected telomeric proteins in PDS-treated over untreated cells.

### LIG3 repairs PDS-damaged telomeres

To address the roles of LIG3 at PDS-induced telomere fragility we depleted LIG3 by RNA interference (Fig. 4A). In untreated control cells, LIG3 depletion did not affect the frequencies of telomere fragility and outsiders (Fig. 4B-D). However, LIG3 depletion reduced the PDS-induced fragility to the levels of untreated cells. In contrast to fragility, the frequency of PDS-induced outsiders increased approximately two-fold upon LIG3-depletion. These results indicate that G4-stabilization by PDS induces telomere cleavage which is repaired in a LIG3-dependent manner. The increase in telomere fragility in response to PDS treatment is due to LIG3-dependent DNA repair. The LIG3-mediated telomeric repair products may give a fragile appearance because of incomplete chromatin condensation subsequent to DNA repair or due to presence of partially single stranded DNA after repair which resists chromatin compaction. We also quantified the frequency of telomere damage induced foci (TIF) (Takai *et al*, 2003) by detection of 53BP1 foci colocalizing with telomeres (Fig. 4E, F). PDS treatment led to an increase in TIF foci, which was further exacerbated upon LIG3 depletion as expected for persistent telomere damage.

**Figure 4.**
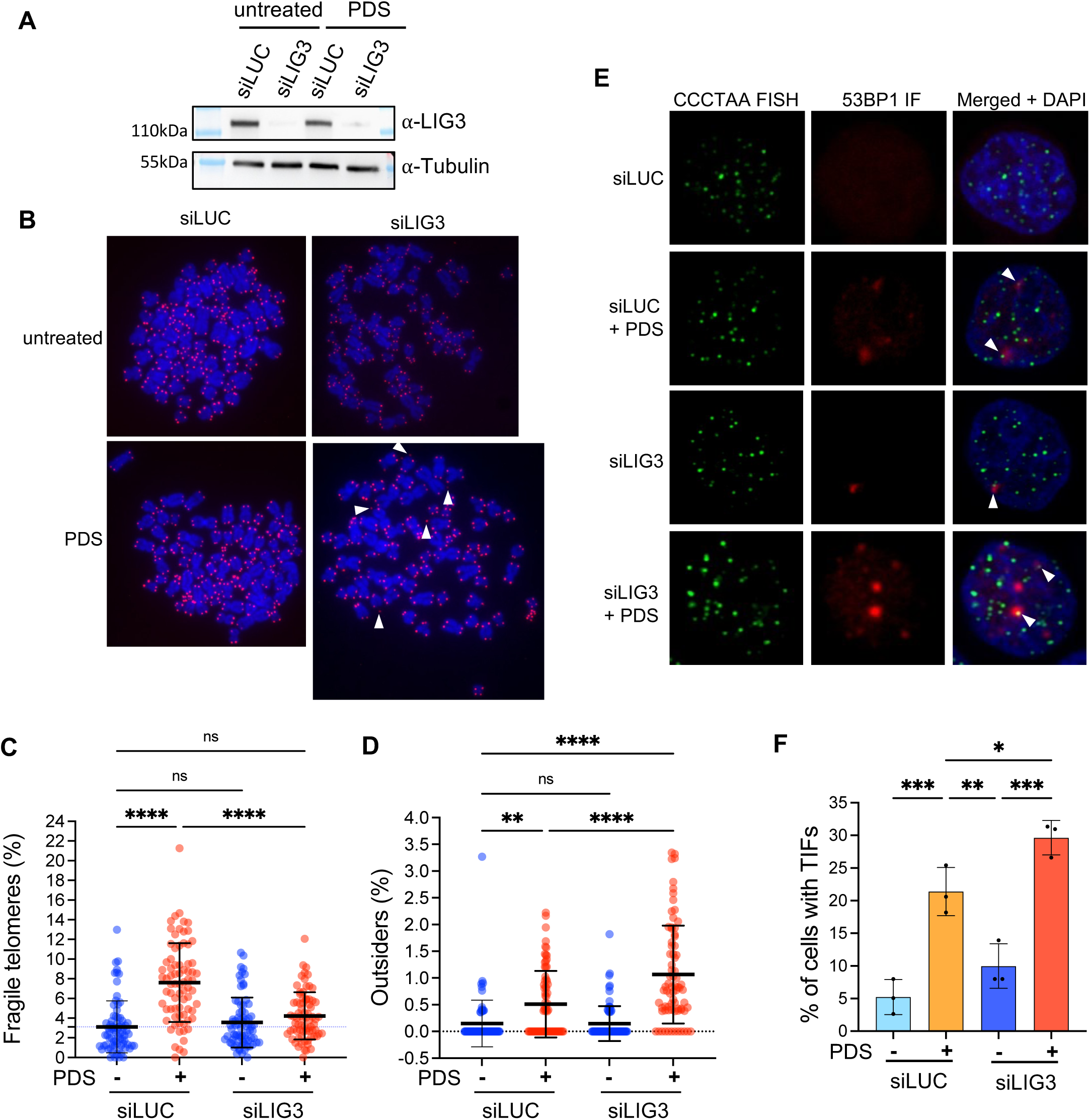
LIG3 repairs PDS-damaged telomeres. (**A**) Western blot analysis of siRNA depletion (siLIG3) of LIG3 in HeLa cells. Tubulin was detected as a loading control. (**B**) Representative telomeric FISH images of metaphase chromosome spreads in presence and absence of LIG3 and upon treatment with PDS. Outsider telomeres are indicated with white arrowheads. Quantification of fragile telomeres (**C**) and outsider telomeres (**D**) upon LIG3 depletion and with or without PDS treatment. 75 metaphase spreads were analyzed per condition, across three independent biological replicates. Horizontal lines and error bars represent mean ± s.d. P-values are calculated using ordinary one-way ANOVA (\*\*\*\**P* < 0.0001, \*\**P* < 0.0016). (**E**) Representative images for detection of 53BP1 at telomeres in presence or absence of PDS and LIG3 depletion as indicated. IF for 53BP1 (red) was combined with telomeric Cy3-[CCCTAA]_3_ FISH (green) and DAPI staining (blue). White arrowheads indicate 53BP1 foci co-localizing with telomeres. (**F**) Quantification of the number of cells with more than one 53BP1 foci co-localizing with telomeres. Bars represent mean ± s.d. of three independent experiments. P-values are calculated using two-way ANOVA (\*\*\**P* = 0.0002, \*\**P* = 0.0043).

### LIG3 depletion suppresses PDS-induced lagging strand telomere fragility while increasing unequal sister chromatid exchange

In order to distinguish strand-specific roles of LIG3 at telomeres, we quantified telomere fragility in cells with >30 kb telomeres in which telomeres were stained with the CO-FISH protocol (Fig. 5A, B). As seen in the CO-FISH analysis of Figure 1, telomere fragility increased with PDS only at lagging strand telomeres (Fig. 5C). Significantly, the increased telomere fragility at lagging telomeres upon PDS treatment was fully suppressed by LIG3 depletion (Fig. 5A, C). At leading strand telomeres, PDS treatment and LIG3-depletion had no significant effects on fragility. Therefore, PDS-mediated telomere damage is repaired by LIG3 specifically at lagging strand telomeres. The CO-FISH experiment also revealed that LIG3-depletion in PDS-treated cells caused a slight but significant increase in the frequency of telomeres in which both sisters had been replicated by leading strand DNA synthesis (Fig. 5B, D). These so-called double-leading telomeres must have occurred due to illegitimate recombination events providing an alternative route of repair in the absence of LIG3 of the PDS-induced DNA double strand breaks.

**Figure 5.**
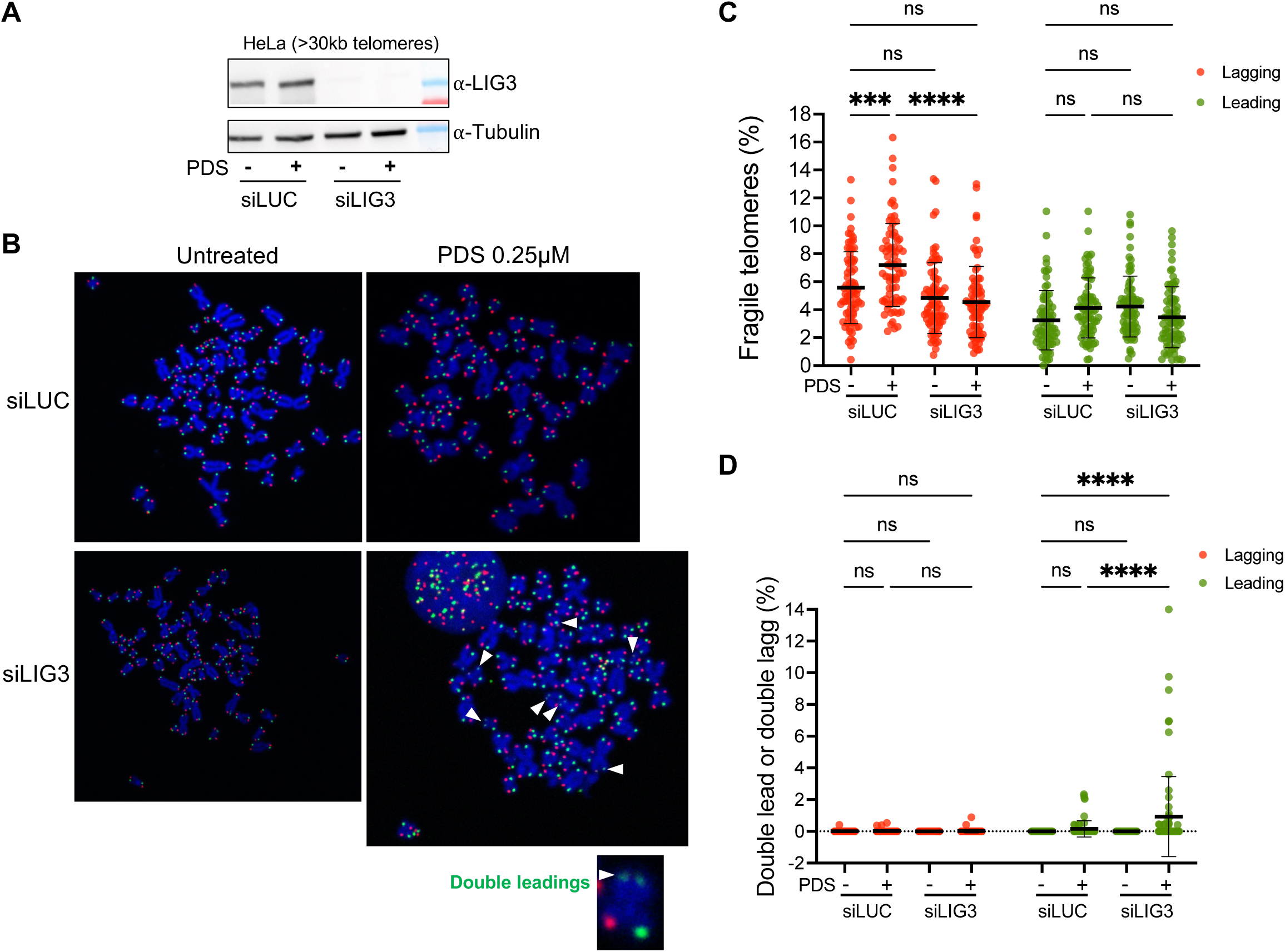
LIG3 depletion in PDS treated cells reduces lagging strand telomere fragility. (**A**) Western blot analysis of siRNA depletion (siLIG3) of LIG3 in HeLa cells with >30 kb telomeres. (**B**) Images of chromosomes with telomere abnormalities analyzed CO-FISH obtained from LIG3-depleted cells that were treated with PDS. Lagging strand telomeres are in red and leading strand telomeres in green. White arrowheads indicate double leading telomeres with fainter green signals at both sister telomeres. (**C**) Quantification of fragile lagging or leading strand telomeres. (**D**) Quantification of chromosomes with double leading or lagging sister telomere signals. 75 metaphase spreads were analyzed per condition, across three independent biological replicates. Horizontal lines and error bars represent mean ± s.d. P-values are calculated using ordinary one-way ANOVA (\*\*\**P* = 0.0004, \*\*\*\**P* < 0.0001).

### PDS-damaged telomeres are repaired by MMEJ

LIG3 participates in several DNA repair pathways including MMEJ, base-excision repair and single strand break repair (Cappelli *et al*, 1997; Simsek *et al*, 2011; Billing & Sfeir, 2025). Since LIG3-depletion in PDS treated cells led to decrease of fragile telomeres we suspected the involvement of MMEJ for repair of PDS-damaged telomeres. We tested this hypothesis by chemically inhibiting or depleting other MMEJ factors than LIG3 by assessing the frequency of telomere fragility and outsiders in presence and absence of PDS (Fig. 6). DNA polymerase theta (Polθ) is an essential component of MMEJ. Chemical inhibition of either the polymerase activity of Polθ with ART558 (Zatreanu *et al*, 2021) or of its helicase activity with AB25583 (Ito *et al*, 2024) abolished the increased telomere fragility induced by PDS (Fig. 6A, B). Similarly, depletion of the APE2 3’-5’ exonuclease and endonuclease suppressed PDS-induced telomere fragility (Fig. 6C, D). APE2 participates in MMEJ as well as in base excision repair (Johnson *et al*, 1998; Simsek *et al*, 2011). Furthermore, co-depletion of LIG3 with inhibition of Polθ or depletion of APE2 revealed that the effects are epistatic (Fig. 6A-D). Overall, the data indicate that MMEJ is responsible for the telomere fragility elicited by PDS.

**Figure 6.**
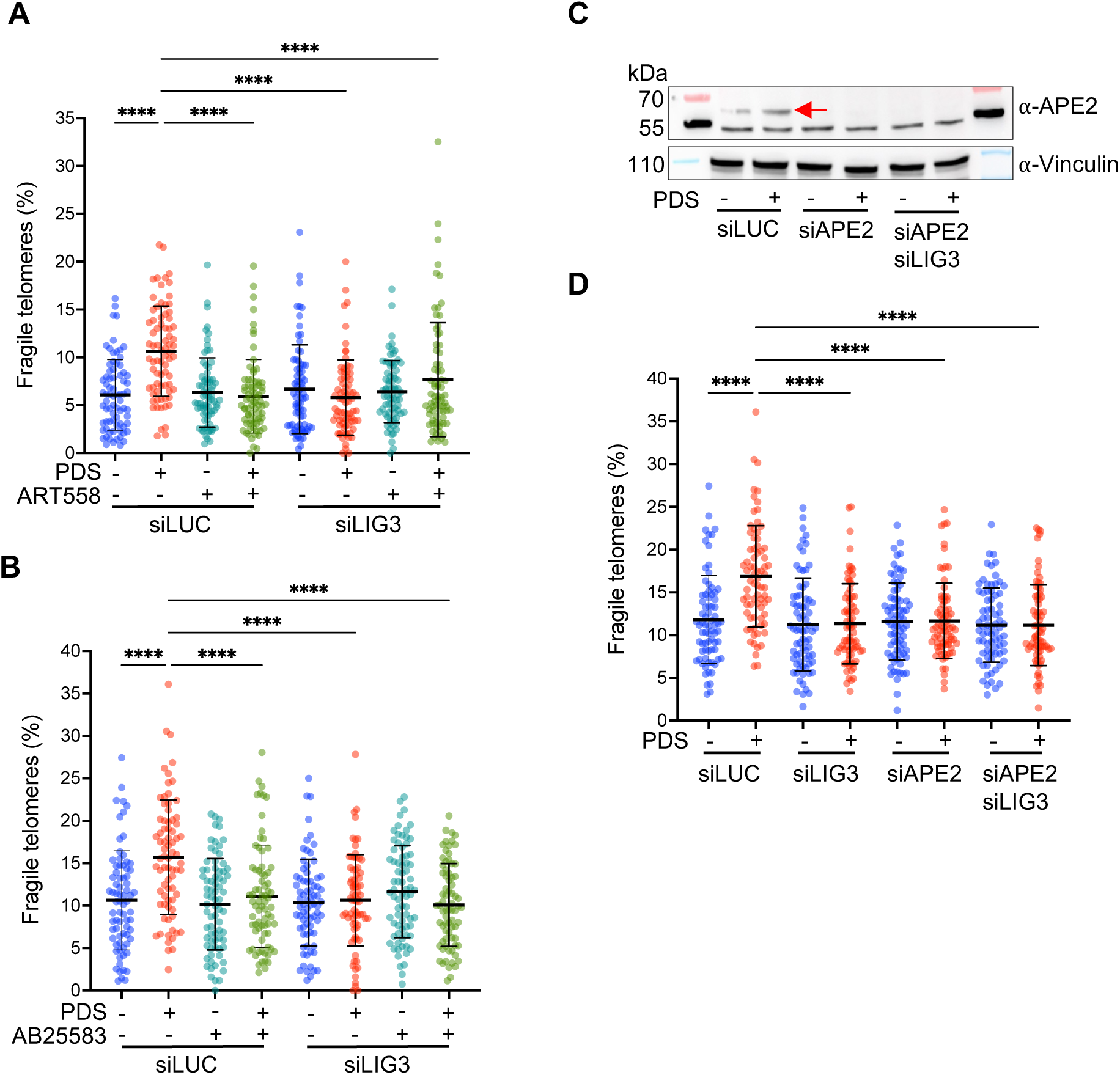
PDS damaged telomeres are repaired by MMEJ. (**A**) Quantification of fragile telomeres after 24 h of treatment of HeLa cells with PDS and POLΣ Polymerase inhibitor (ART558) as indicated. (**B**) Quantification of fragile telomeres after 24 h of treatment of HeLa cells with PDS and POLΣ ATPase inhibitor (AB25595) as indicated. (**C**) Western blot analysis of siRNA depletion (siAPE2) of APE2 in HeLa cells. Red arrow indicates APE2 (**D**) Quantification of fragile telomeres after APEX2 depletion using siRNA and PDS treatment of HeLa cells as indicated. 75 metaphase spreads were analyzed per condition, across three independent biological replicates. Horizontal lines and error bars represent mean ± s.d. P-values are calculated using two-way ANOVA (\*\*\*\**P* < 0.0001).

### PDS-induced telomere cleavage requires DNA2 and FEN1

As demonstrated in Fig.4, PDS-treatment in LIG3-depleted cells caused robust accumulation of telomere cleavage and outsider telomeres. In order to identify the enzymes involved in telomeres cleavage of PDS-treated cells, we depleted candidate nucleases and quantified their effects on outsider frequency (Fig. EV3). siRNA-mediated depletion of either FEN1 (Fig. EV3A) or DNA2 (Fig. EV3B) completely suppressed the outsider phenotype of PDS-treated LIG3-depleted cells (Fig. EV3C). Thus, both enzymes are required for telomere cleavage and for subsequent initiation of MMEJ-mediated telomere repair.

## Discussion

PDS and G4 structures have been implicated to induce telomere damage by various means including interference with the binding of shelterin components and the CST complex, interference with DNA replication and telomerase inhibition (Johnson *et al*, 2024; Rodriguez *et al*, 2012; Zimmer *et al*, 2016; Groelly *et al*, 2022; Crabbe *et al*, 2004; Bryan, 2020; Varshney *et al*, 2020). Here, we demonstrate that PDS acts already before telomerase-mediated telomere elongation during the semiconservative replication of telomeric DNA. Our analysis suggests a three-step model (Fig. 7). First, PDS interferes specifically with lagging strand DNA synthesis of telomeres indicating binding to the lagging strand G-rich parental template, stabilizing G4-structures and sequestering it from replicative polymerases. PDS may favor G4-structures on the parental DNA during replication upon unwinding by the CMG replicative helicase. Alternatively, PDS may stabilize G4-structures ahead of the replication fork possibly binding to the displaced G-rich DNA strand present in R-loop structures induced by the TERRA long noncoding RNA (Kyriacou & Lingner, 2024). The impediment at the replication fork leads to ATR signaling as evidenced by the accumulation of P-Ser33 RPA32 foci as well as replication fork stalling as seen in the telomeric DNA fiber analysis. Second, Flap-endonuclease FEN1 in conjunction with the nuclease-helicase DNA2 cleave the G4-containing DNA removing the G4 structures. Consistent with our results, DNA2 was shown to cleave G4-containing DNA *in vitro* (Lin *et al*, 2013) as well as *in vivo* (Fernandez *et al*, 2025). Our data indicate that in addition to DNA2, FEN1 is required for cleaving the G4-containing lagging strand telomeres. Thus, the two enzymes which cooperate during Okazaki-fragment processing appear to require each other for the cleavage reaction. Third, MMEJ ligates the broken telomeric DNA fragments. Telomere healing is dependent on the helicase and polymerase activities of Polθ, APE2 3’-5’ exonuclease and endonuclease, and LIG3. The final MMEJ-repaired telomeres leave behind scars in the form of fragile telomeres, with no apparent effect on telomere function and cell growth.

**Figure 7.**
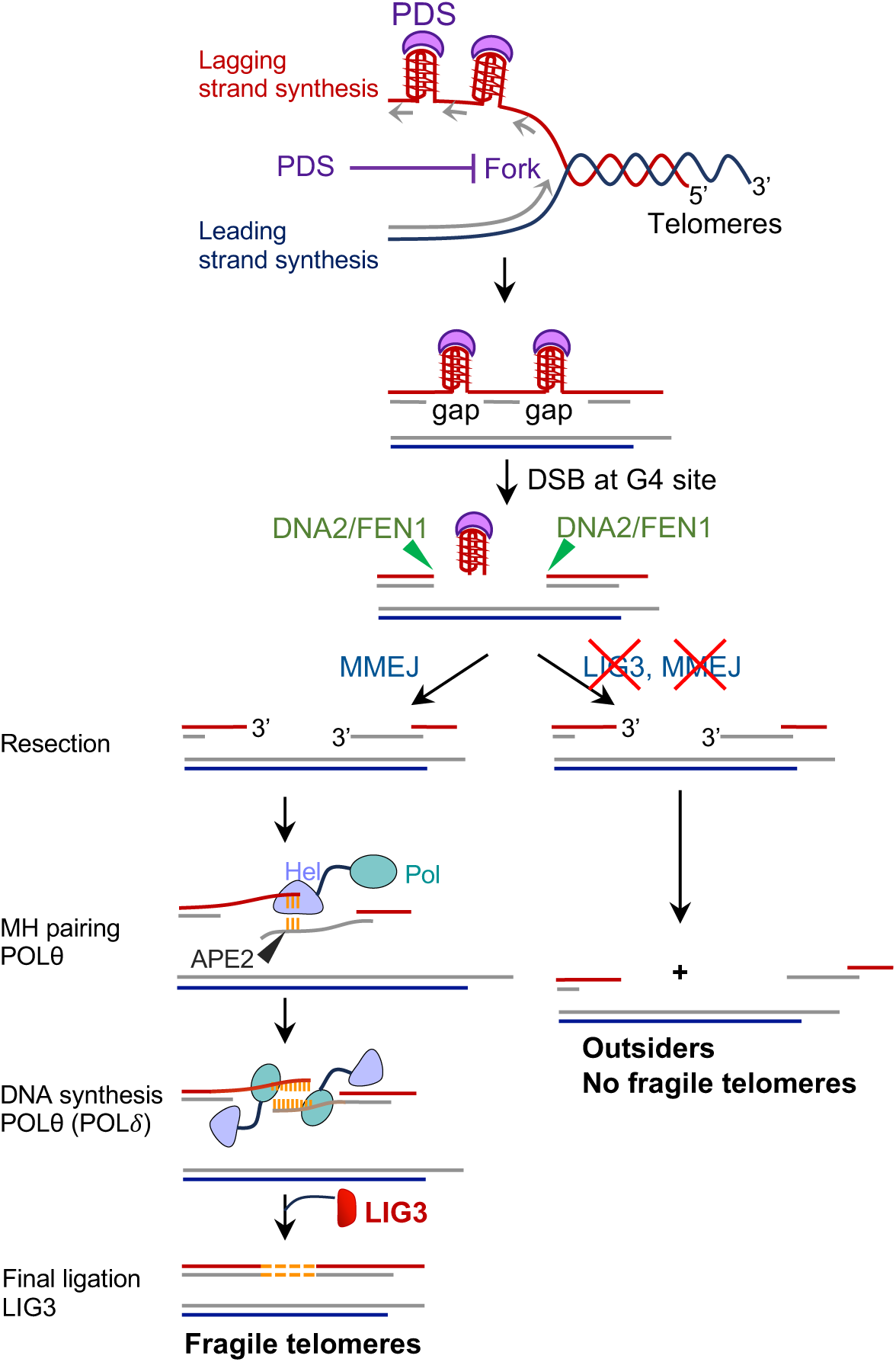
Model. PDS binds during semiconservative DNA replication to G4 structures of the parental G-rich telomeric strand that serves as a template for discontinuous lagging strand DNA synthesis. The PDS-trapped G4 structures cannot be used as a template leading to presence of single stranded DNA gaps which may elicit ATR signaling and RPA phosphorylation. PDS also triggers replication fork arrest. To overcome this hurdle, G4-structures are removed by DNA cleavage involving DNA2 and FEN1. The broken telomeric DNA fragments are stitched together by MMEJ giving rise to telomere fragility. Upon inhibition of MMEJ in PDS-treated cells, the broken telomeric DNA is not repaired and outsider telomeres accumulate.

Our data demonstrate that concomitant treatment of cancer cells with G4-stabilizing drugs and MMEJ-inhibitors induce telomere cleavage thus inducing sudden telomere loss and strongly enhancing the damage at telomeres. MMEJ-deficiency has been recently reported for Ewing sarcomas (Asada *et al*, 2025) suggesting that these cancers may be particularly sensitive to PDS treatment. PDS as well as other G4-stabilizing drugs have also been considered as therapeutic agents to target telomeres in BRCA1 and BRCA2-mutant cancer cells that are defective in homology-directed repair (HDR) (Zimmer *et al*, 2016; Groelly *et al*, 2022; Keahi *et al*, 2025) and clinical trials with G-stabilizers for HDR-deficient tumors have been launched (Hilton *et al*, 2022).

Finally, telomerase inhibition can lead to slow telomere shortening killing cancer cells in a delayed fashion after longer periods of growth that may be needed to obtain critically short telomeres. Our data suggest that a combined treatment with G4-stabilization and MMEJ-inhibition may synergize with future anti-telomerase therapies to accelerate their effects on cancers with long telomeres.

## Methods

### Cell culture

HeLa cells with an average telomere length of 10kb and HeLa cells with an average telomere length of more than 30kb (Cristofari & Lingner, 2006) were cultured in Dulbecco’s Modified Eagle Medium (DMEM) supplemented with 10% fetal bovine serum (FBS) (Clontech) and 100U/ml of Penicillin/Streptomycin (Gibco) at 37°C in presence of 5% (v/v) CO_2_. Suspension HEK293E cells were cultured in EX-CELL 293 Serum-Free Medium, supplemented with 4 mM GlutaMAX with 5% (v/v) CO_2_, at 37°C, with constant agitation. Stable cell lines expressing endogenous 3xflag-TRF1 and 3xflag-TRF2 (heterozygous) were described before (Lin *et al*, 2021).

### siRNA transfection

siRNAs were transfected using calcium phosphate. 200,000 cells were seeded in six-well plates one day before siRNA transfection. A 100 μL transfection mix was prepared, consisting of 125 mM CaCl₂, 110 nM siRNA, and 1×HBSS (pH 7.4) containing 50 mM HEPES, 280 mM NaCl, 1.19 mM Na₂HPO₄•2H₂O, and 10 mM KCl. The mix was incubated at RT for 1 min before being added to the cells in 1 mL of antibiotic-free DMEM supplemented with 10% FBS. Cells were treated with 0.12 or 0.25 µM PDS for 24 h before harvesting.

### Western blotting

Cells were harvested by trypsinization, washed with 2× Laemmli buffer and resuspended to a final concentration of 10,000 cells/µl of Laemmli buffer. Samples were incubated at 95°C for 5min. Proteins were separated by SDS-PAGE and transferred onto a 0.2 µm nitrocellulose membrane (GE Healthcare). Membranes were blocked with blocking solution (5% milk in 1×PBS with 0.1% Tween-20 (PBST)) for 1 h at room temperature (RT) and incubated with primary antibodies (see “Reagent and Tools Table”) at 4°C overnight. Membranes were washed three times for 5min with PBST. Horseradish Peroxidase-conjugated secondary antibodies (see “Reagent and Tools Table”) in combination with ECL spray (Advansta) were used to reveal the signal on a Fusion FX (Vilber) detector.

### Immunofluorescence and telomeric fluorescence in situ hybridization (IF-FISH)

HeLa cells were cultured on round coverslips and treated with 0.12µM PDS for 24 h, washed twice with PBS, and fixed in 4% paraformaldehyde for 10 minutes. After fixation, cells were permeabilized with a detergent solution (0.1% Triton X-100, 0.02% SDS in PBS) and blocked with 2% BSA in PBS for 10 minutes. Primary antibody (anti-53BP1, 1:2,000) was diluted in a blocking solution (10% normal goat serum, 2% BSA in PBS) and incubated overnight at 4°C in a humid chamber. After multiple washes, secondary antibody (anti-rabbit Alexa Fluor 633, 1:800) was applied for 30 minutes. Coverslips were fixed again in 4% paraformaldehyde, washed, and dehydrated with a graded ethanol series (70%, 95%, and 100%). For pRPA IF-FISH, HeLa cells cultured on round coverslips were pre-extracted with CSK buffer (10 mM PIPES pH 7.0, 100 mM NaCl, 300 mM sucrose, 3 mM MgCl₂, 0.5% Triton X-100). Cells were fixed with 4% formaldehyde for 10 min, permeabilized with CSK buffer and blocked in PBST containing 0.5% BSA for 1 h. Primary antibody (anti-pSer33 RPA32, 1:2000) was added overnight at 4°C in a humid chamber. After three washes with PBST, secondary antibody (anti-rabbit Alexa fluor 633nm, 1:1,000) was added for 50 min. Cells were washed three times for 10 min with PBST. Coverslips were fixed again with 4% formaldehyde and dehydrated. FISH staining was performed as previously described (Fernandes & Lingner, 2023). Coverslips were mounted with Vectashield, and images were acquired with a Leica SP8 confocal microscope equipped with a 63 ×/ 1.40 oil objective.

### Telomeric FISH on metaphase chromosomes

Telomeric FISH was performed as previously described (Fernandes & Lingner, 2023). Briefly, cells were treated with 0.05 μg/mL demecolcine for 2 hours before being harvested and subjected to hypotonic treatment (0.056 M KCl) at 37°C for 7 minutes. After fixation in cold methanol/acetic acid (3:1) overnight, cells were spread onto slides and processed for FISH staining, using a Cy3-TeloC PNA probe (100 nM). Images were acquired using an Upright Zeiss Axioplan equipped with a 100x/1.40 oil objective.

### Chromosome Orientation (CO)-FISH on metaphase chromosomes

CO-FISH was performed as previously described (Fernandes & Lingner, 2023). Briefly, HeLa cells with >30kb telomeres were incubated with a mix of BrdU/BrdC (3:1) at a final concentration of 10 μM for 15 hours and treated with 0.1 μg/mL demecolcine for 2 hours. After hypotonic treatment, cells were fixed and spread onto glass slides. After rehydration, cells were treated with RNaseA, stained with Hoechst 33258, and exposed to UV light. Following digestion with exonuclease III and paraformaldehyde fixation, hybridization was performed using TYE563-TeloC LNA and 6-FAM-TeloG LNA probes. Post-hybridization washes were performed, and slides were mounted with ProLong Diamond Antifade mountant. Images were acquired using a Leica SP8 confocal microscope equipped with a 63 ×/ 1.40 oil objective.

### 2-step QTIP

Telomeric chromatin precipitation was performed as previously described (Glousker *et al*, 2020) with some modifications. One billion of HEK293E cells expressing endogenous 3xflag-TRF1 and 3xflag-TRF2 were used per IP reaction. HEK293E cells were treated with 15 µM PDS for 24 h. After chromatin enrichment, sonicated extracts were dialyzed. The first immunoprecipitation (IP) was performed using either Anti-flag M2 affinity Agarose Gel or mouse IgG coupled to Sepharose beads. Purified proteins were eluted with flag peptide. A second IP was performed using home-made affinity-purified anti-TRF1 and anti-TRF2 antibodies coupled to protein G Sepharose beads. After the second step of purification, telomeric chromatin was eluted with ammonium hydroxide. Purification efficiency was evaluated by quantification of telomeric DNA and Alu-repeat DNA signals on Dot blots (Fig. EV2B). Fractions were pooled and flash-frozen. Samples were then lyophilized for mass spectrometry analysis. Lyophilized samples were digested with a Trypsin/LysC mix. After digestion samples were pooled in two Tandem Mass Tag (TMT)-mixes. Four replicates of untreated and PDS treated samples were combined in a height-plex TMT experiment and negative IgG controls in a six-plex TMT experiment. No fractionation was applied to the samples.

527 proteins were identified in IP samples and 81 proteins in IgG controls. Common proteins detected in both IgG fractions and 2-step QTIP fractions were trimmed leaving 501 candidates (Fig. EV2C, Table EV1). To determine significant changes between untreated and PDS treated conditions, differential protein expression analysis was performed using the R Bioconductor package LIMMA followed by Benjamini-Hochberg multiple-testing correction method.

### Proximity ligation assay (PLA)

HeLa cells with >30kb telomeres were cultured on round coverslips and treated with either 0.12 µM or 10 µM of PDS for 24 h. After CSK pre-extraction, cells were fixed with 4% formaldehyde for 15 min. Coverslips were then washed with PBS, permeabilized with 0.5% Triton X-100 in PBS for 10 min at RT and washed again with PBS. Cells were incubated with RNase A (10 µg/ml) for 1 h at 37°C. After three washes with PBS, cells were blocked with 3% BSA in PBS for 1 h. Primary antibodies (anti-G4 (1H6) and anti-TRF2, 1:500) were incubated overnight at 4°C. After three washes with PBST, cells were incubated with anti-mouse minus and anti-rabbit plus PLA probes at 37°C for 1 h. Cells were washed with PLA buffer A (10 mM Tris pH 7.4, 150 mM NaCl, 0.05% Tween 20) and incubated with ligation buffer containing ligase at 37°C for 30 min, then washed with buffer A and incubated with amplification buffer with polymerase at 37°C for 100 min. Cells were washed twice with PLA buffer B (200 mM Tris pH 7.5, 100 mM NaCl) and then three times with PBST buffer containing DAPI. Coverslips were mounted with Vectashield, and images were acquired using a Leica SP8 confocal microscope equipped with a 63 ×/ 1.40 oil objective.

### Single molecule analysis of replicated DNA (SMARD)

SMARD was performed as previously described (Norio & Schildkraut, 2001) with some modifications. HeLa cells with >30kb telomeres were sequentially labeled at 37°C with 5-chloro-2’-deoxyuridine (CldU, 100 µM) for 15 min and 5-iodo-2’-deoxyuridine (IdU, 100 µM) for 15 min with three PBS washes in between. Where indicated, 0.25 µM PDS was added together with IdU. Labelling was repeated three times with intermittent 2 h incubations in label-free medium. One million labeled cells were embedded in 0.7 % low melting point agarose. After cell lysis in 0.5 M EDTA, 1% N-lauryl sarcosine, 0.2 mg/ml proteinase K overnight at 50°C, restriction digestion was performed using HinfI and RsaI to fragment non-telomeric DNA. Telomeric DNA was separated on 0.7% low melting-point agarose using pulse field gel electrophoresis. Telomeric DNA was extracted from gel slices by digestion with ß-agarase I overnight at 42°C. 1 µg of purified telomeric DNA was spread on 3-aminopropylthriethoxysilate-coated glass slides and incubated with 0.1% ß-mercaptoethanol in methanol for 10 min at RT. Following denaturation with denaturation buffer (0.1 N NaOH, 0.1% ß-mercaptoethanol in 70% ethanol) for 10 minutes, the slides were fixed with 0.5% glutaraldehyde in denaturation buffer for 5 min at RT. Slides were dehydrated with sequential incubation with 70%, 95% and 100% ethanol and incubated again in 0.1% ß-mercaptoethanol in methanol for 1 h. Telomeric DNA was identified by hybridization with a TeloC-Biotin conjugated PNA probe (500 nM) at 37°C overnight. Slides were incubated with 40% formamide in 4X SSC at 40°C for 5 min. Following 4 washes with 0.1% IGEPAL CA-630 in 4X SSC, slides were incubated in blocking solution (3% BSA, 0.03% IGEPAL CA-630 in PBS) for 1h at RT. The biotinylated telomeric probe was detected by incubation with NeutrAvidin Alexa fluor 350 antibody (1:500 in blocking solution). Telomeric signals were enhanced by alternating rounds of incubation with biotinylated anti-avidin and NeutrAvidin antibodies. Rat anti-BrdU antibody (1:500) and mouse anti-BrdU antibody (1:200) were used to detect CldU and IdU tracks respectively. Anti-rat Cy3 (1:150) and anti-mouse Alexa fluor 488 were used as secondary antibodies (1:300) (see “Reagent and Tools Table”). Slides were mounted with ProLong Diamond Antifade mounting media and images were acquired using the Leica SP8 confocal microscope equipped with a 63 ×/ 1.40 oil objective.

### Reagents and Tools Table

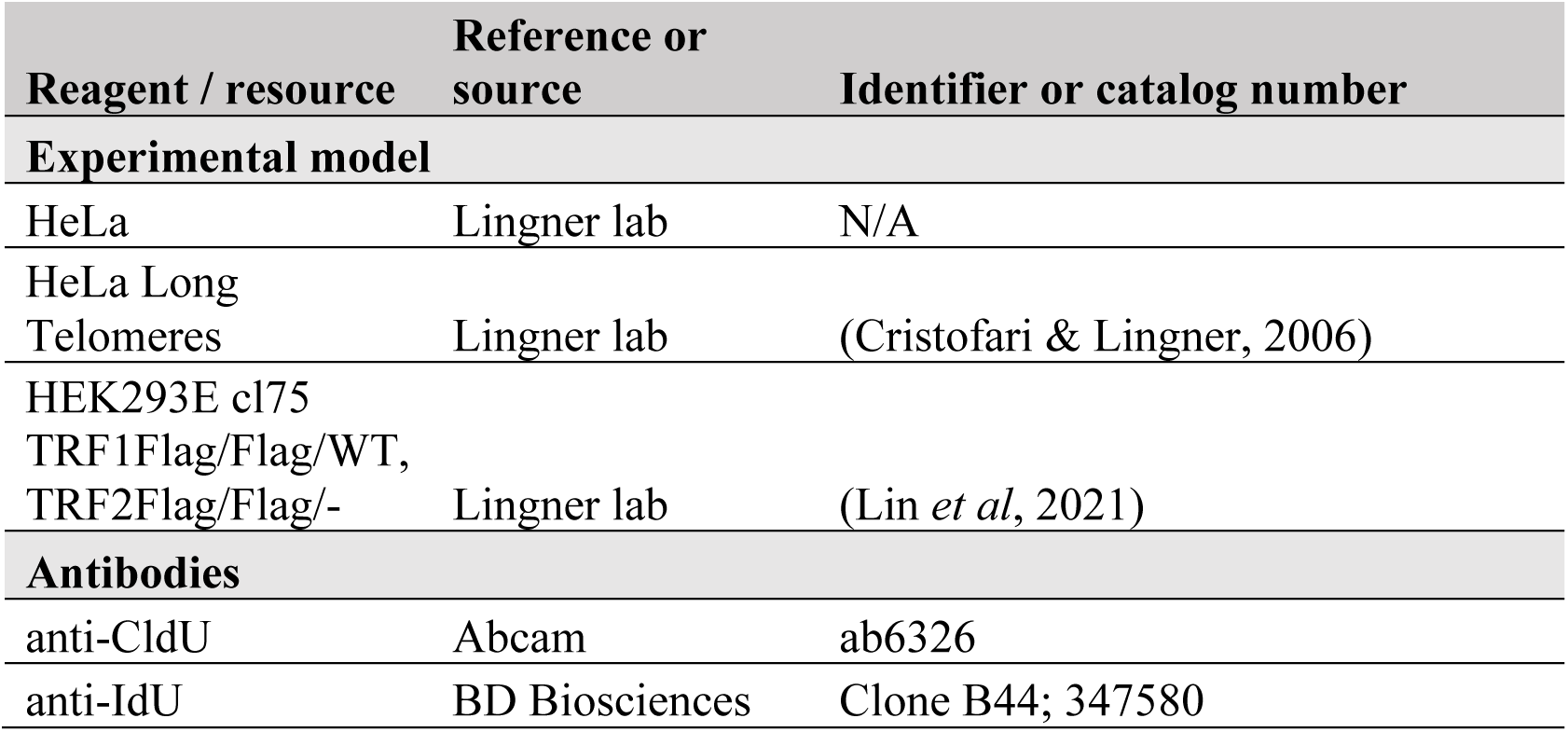
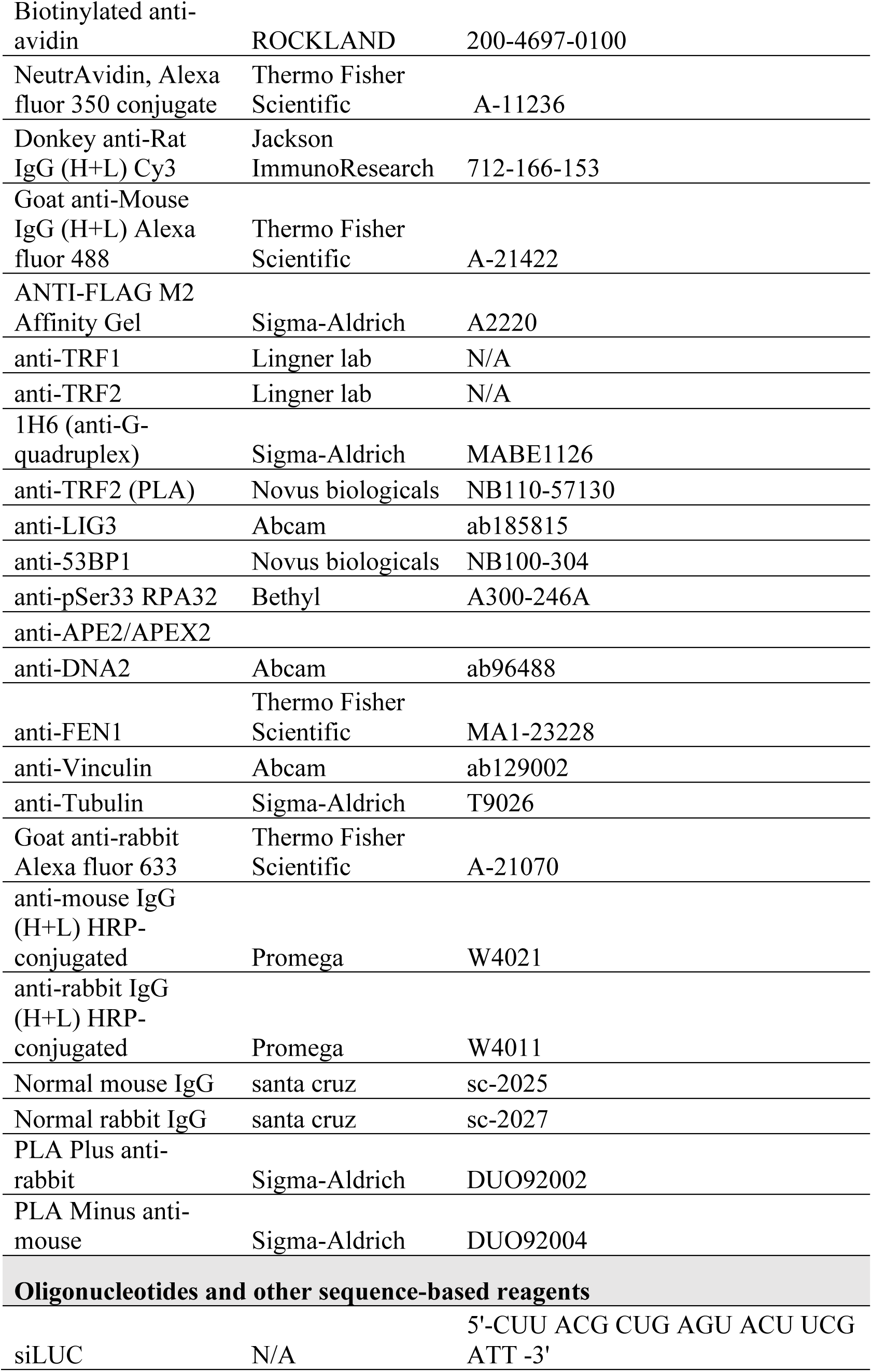
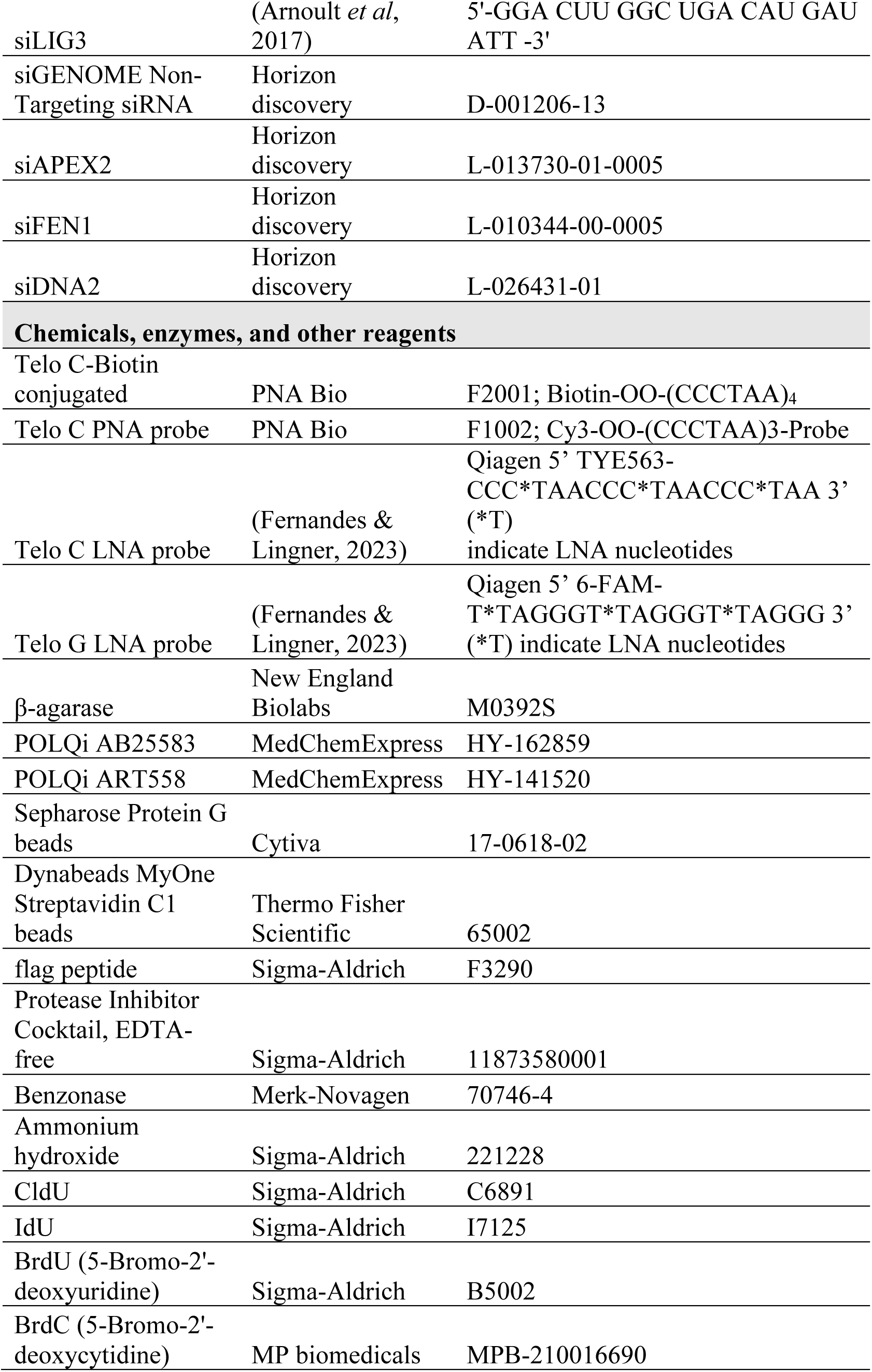
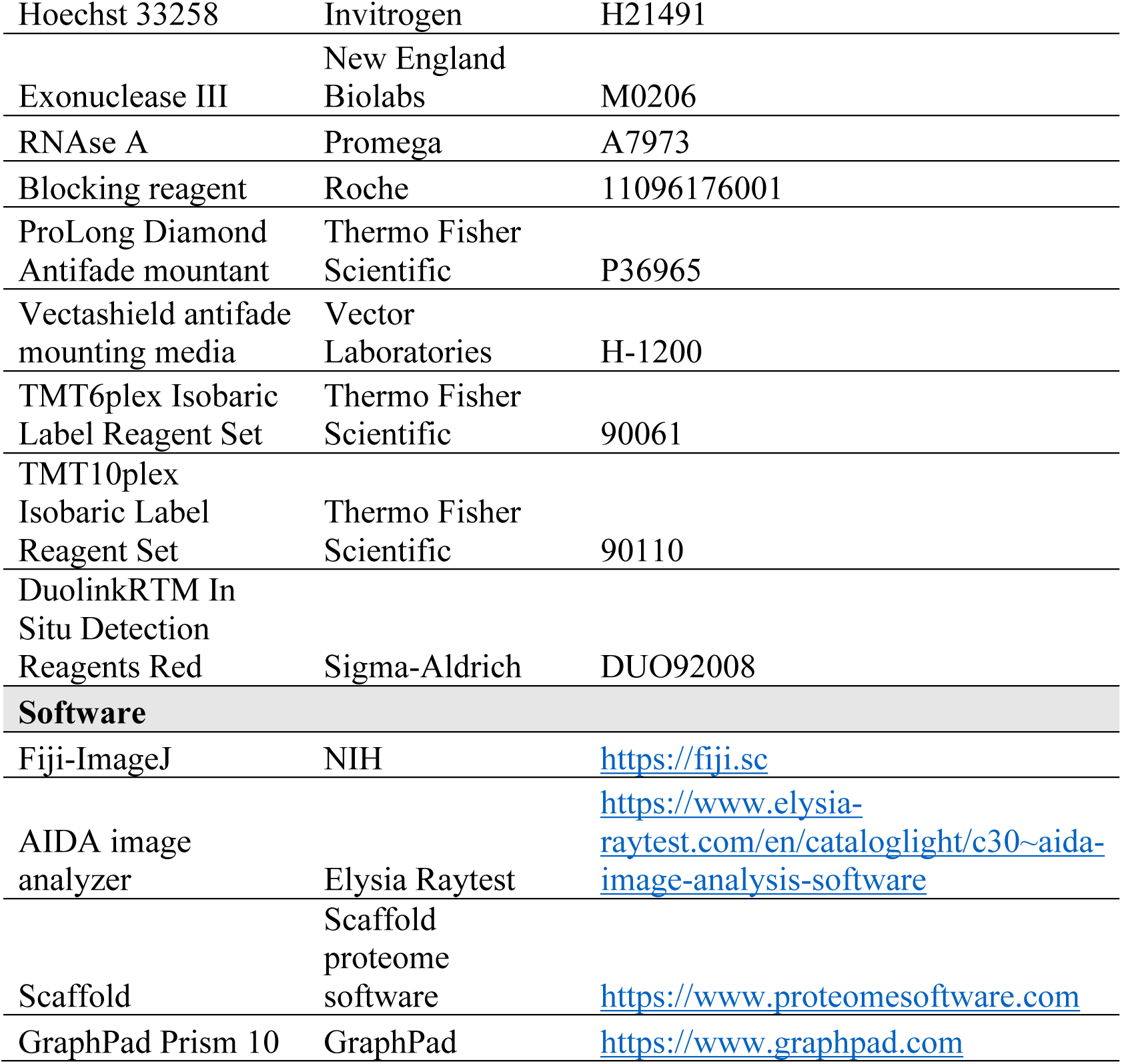

## Acknowledgements

We thank Galina Glousker, Eftychia Kyriacou and Misun Jung for comments on the manuscript. We acknowledge the support provided by the Proteomics Core Facility at the School of Life Sciences of EPFL. This work was supported by the Swiss National Science Foundation (SNSF) [310030_214833] and the SNSF-funded National Centre of Competence in Research RNA and Disease Network [205601].

## Author contributions

**Samah Matmati**: Conceptualization; Methodology; Investigation; Formal analysis; Data curation; Writing; **Satyajeet Rao**: Methodology; Investigation; Formal analysis; **Joachim Lingner**: Conceptualization; Supervision; Funding acquisition; Writing.

## Expanded View Figures

**Figure EV1.**
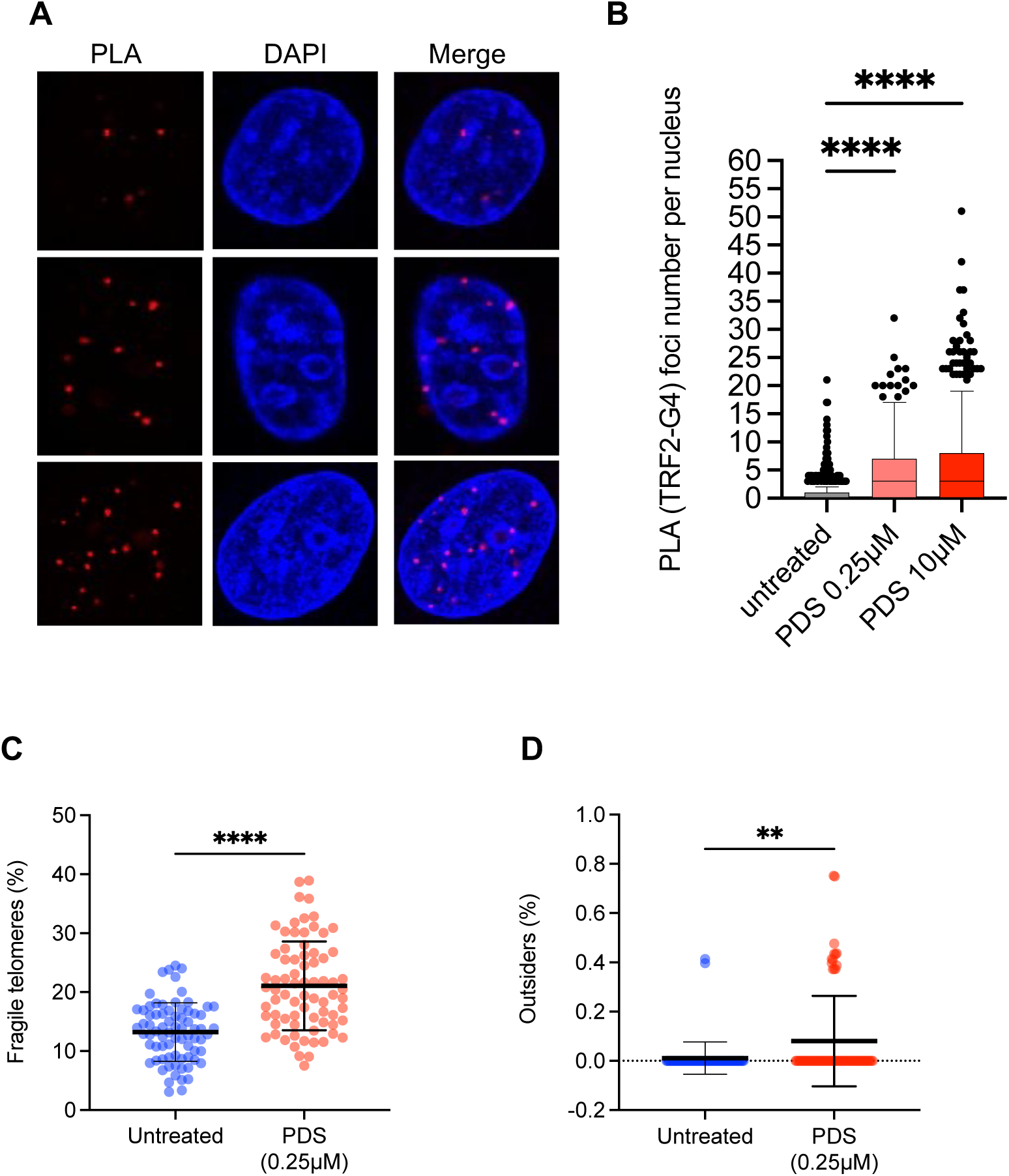
Effects of PDS treatment in HeLa cells, related to Fig 1. and 2. (**A**) Proximity ligation assay (PLA) for detection of telomeric G4. HeLa cells (>30 kb telomeres) were analyzed by PLA using anti-G4 (1H6) and anti-TRF2 antibodies. Representative images of PLA foci (red) and DAPI (blue). More than 600 nuclei were analyzed per condition, across three independent biological replicates. (**B**) Tukey boxplot analysis of PLA foci per nucleus in untreated and PDS-treated cells (0.25 µM or 10 µM as indicated). Background PLA signals obtained from single anti-TRF2 antibody were subtracted. Horizontal lines represent the interquartile range. P-values are calculated from ordinary one-way ANOVA (\*\*\*\**P* < 0.0001). (**C**), (**D**) Quantification of fragile telomeres (C) and outsiders (D) obtained upon telomeric FISH analysis of metaphase chromosome spreads of HeLa cells (with >30 kb average telomere length). 0.25 µM PDS treatment lasted for 24 h. 75 metaphase spreads were analyzed per condition, across three independent biological replicates. Horizontal lines and error bars represent mean ± s.d. P-values are calculated from Mann Whitney test (\*\*\*\**P* < 0.0001, \*\**P* = 0.0025).

**Figure EV2.**
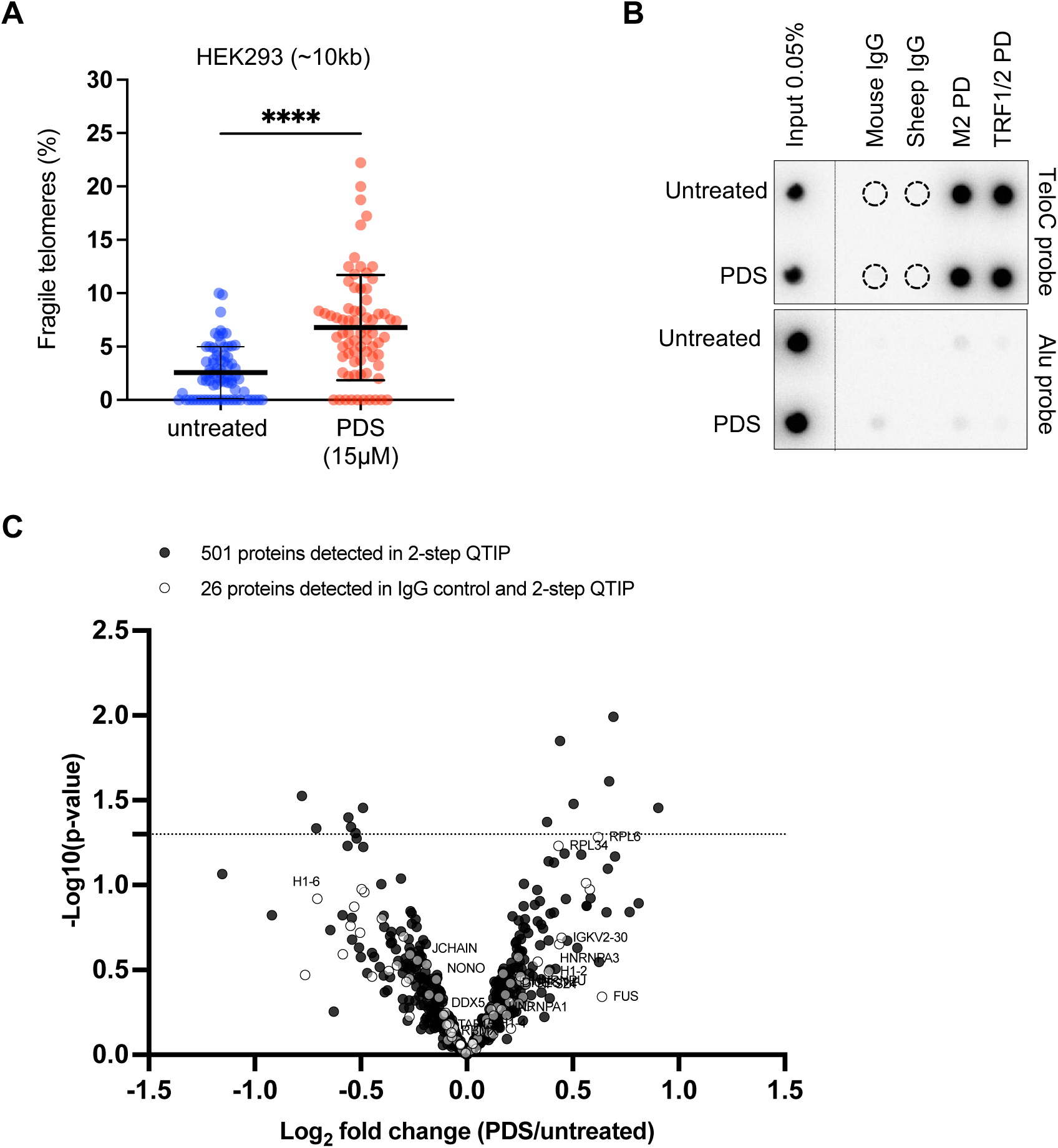
2-step QTIP experiment, related to Fig. 3. (**A**) Quantification of fragile telomeres by telomeric FISH analysis of metaphase chromosome spreads of HEK293E suspension cells treated with 15 µM PDS during 24 h, as indicated. 75 metaphase spreads were analyzed per condition, across three independent biological replicates. Horizontal lines and error bars represent mean ± s.d. P-values are calculated from Mann Whitney test (\*\*\*\**P* < 0.0001). (**B**) Telomeric and Alu-repeat DNA quantification by dotblot analysis during the two steps of 2-step QTIP. DNA in fractions after M2 purification and TRF1/TRF2 purification was detected using ^32^P-labeled C-rich telomeric and ^32^P-labeled Alu probes. IgG negative control fractions were also analyzed. (**C**) Volcano plot representing differential protein expression in untreated versus PDS treated samples in IgG fractions and 2-step QTIP fractions. White dots represent 26 proteins commonly detected in both IgG and 2-step QTIP fractions. Common proteins were eliminated in the final analysis.

**Figure EV3.**
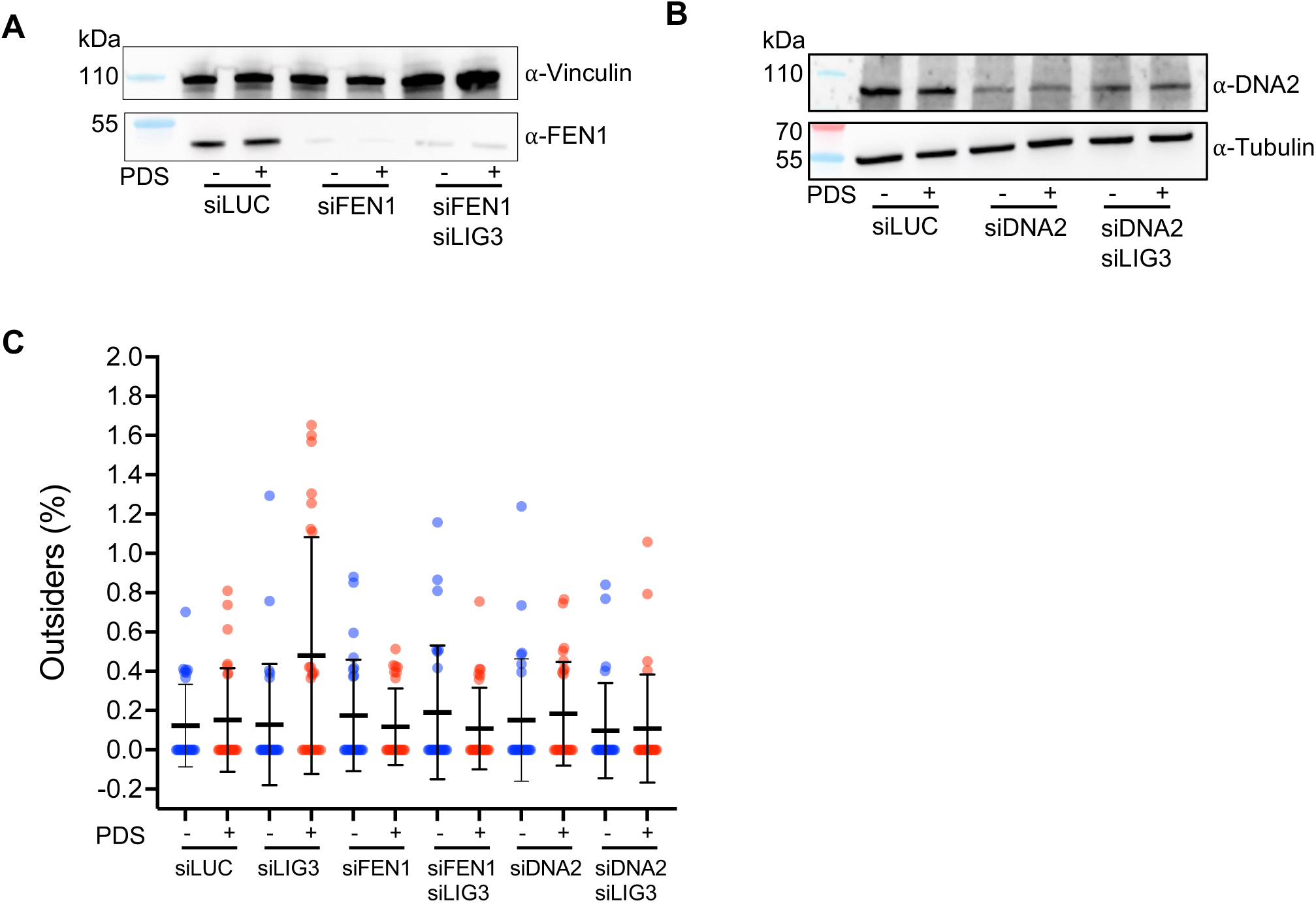
PDS damaged telomeres are processed by DNA2, FEN1 and MMEJ factors, related to Fig. 6. (**A-C**) Western blot analyses of APE2, FEN1 and DNA2 upon siRNA transfections. (**D**) Quantification of outsider telomeres upon 0.12 µM PDS treatment for 24 h in cells depleted for LIG3, FEN1 and DNA2 as indicated.

